# Fast Inference of Individual Admixture Coefficients Using Geographic Data

**DOI:** 10.1101/080291

**Authors:** Kevin Caye, Flora Jay, Olivier Michel, Olivier François

## Abstract

Accurately evaluating the distribution of genetic ancestry across geographic space is one of the main questions addressed by evolutionary biologists. This question has been commonly addressed through the application of Bayesian estimation programs allowing their users to estimate individual admixture proportions and allele frequencies among putative ancestral populations. Following the explosion of high-throughput sequencing technologies, several algorithms have been proposed to cope with computational burden generated by the massive data in those studies. In this context, incorporating geographic proximity in ancestry estimation algorithms is an open statistical and computational challenge. In this study, we introduce new algorithms that use geographic information to estimate ancestry proportions and ancestral genotype frequencies from population genetic data. Our algorithms combine matrix factorization methods and spatial statistics to provide estimates of ancestry matrices based on least-squares approximation. We demonstrate the benefit of using spatial algorithms through extensive computer simulations, and we provide an example of application of our new algorithms to a set of spatially referenced samples for the plant species *Arabidopsis thaliana*. Without loss of statistical accuracy, the new algorithms exhibit runtimes that are much shorter than those observed for previously developed spatial methods. Our algorithms are implemented in the `R package`, `tess3r`.

## 1. Introduction

High-throughput sequencing technologies have enabled studies of genetic ancestry for model and non-model species at an unprecedented pace. In this context, ancestry estimation algorithms are important for demographic analysis, medical genetics including genome-wide association studies, conservation and landscape genetics (Pritchard et al., 2000; Tang et al., 2005; Schraiber et al., 2015; Segelbacher et al., 2010; François et al., 2016). With increasingly large data sets, Bayesian approaches to the inference of population structure, exemplified by the computer program `structure` (Pritchard et al., 2000), have been replaced by approximate algorithms that run several orders faster than the original version (Tang et al., 2005; Alexander et al., 2011; Frichot et al., 2014; Raj et al., 2014). Considering *K* ancestral populations or genetic clusters, those algorithms estimate ancestry coefficients following two main directions: model-based and model-free approaches. In model-based approaches, a likelihood function is defined for the matrix of ancestry coefficients, and estimation is performed by maximizing the log-likelihood function. For `structure` and related models, model assumptions include linkage equilibrium and Hardy-Weinberg equilibrium in ancestral populations. The first approximation to the original algorithm was based on an expectation-minimization algorithm (Tang et al., 2005), and more recent likelihood algorithms are implemented in the programs `admixture` and `faststructure` (Alexander et al., 2011; Raj et al., 2014). In model-free approaches, ancestry coefficients are estimated by using least-squares methods or factor analysis. Model-free methods make no assumptions about the biological processes that have generated the data. To estimate ancestry matrices, Engelhardt et al. (2010) proposed to use sparse factor analysis, Frichot et al. (2014) used sparse non-negative matrix factorization algorithms, and Popescu et al. (2014) used kernel-principal component analysis. Least-squares methods accurately reproduce the results of likelihood approaches under the model assumptions of those methods. In addition, model-free methods provide approaches that are valid when the assumptions of likelihood approaches are not met (Frichot et al., 2014). Model-free methods are generally faster than model-based methods.

Among model-based approaches to ancestry estimation, an important class of methods have improved the Bayesian model of `structure` by incorporating geographic data through spatially informative prior distributions (Chen et al., 2007; Corander et al., 2008). Under isolation-by-distance patterns (Wright, 1943; Malécot, 1948), spatial algorithms provide more robust estimates of population structure than non-spatial algorithms which can lead to biased estimates of the number of clusters (Durand et al., 2009). Some Bayesian methods are based on Markov chain Monte Carlo algorithms which are computer-intensive (François et al., 2010). Recent efforts to improve the inference of ancestral relationships in a geographical context have mainly focused on the localization of recent ancestors (Baran et al., 2013; Lao et al., 2014; Yang et al., 2014; Bhaskar et al., 2017; Ranñola et al., 2014). In these applications, spatial information is used in a predictive framework that assigns ancestors to putative geographic origins. While fast geographic estimation of individual ancestry proportions has been proposed previously (Caye et al., 2016; Bradburd et al., 2016), there is a growing need to develop individual ancestry estimation algorithms that reduce computational cost in a geographically explicit framework.

In this study, we present two new algorithms for the estimation of ancestry matrices based on geographic and genetic data. The new algorithms solve a least squares optimization problem as defined by Caye et al. (2016), based on Alternating Quadratic Programming (AQP) and Alternating Projected Least Squares (APLS). While AQP algorithms have a well-established theoretical background (Bertsekas, 1995), this is not the case of APLS algorithms. Using coalescent simulations, we provide evidence that the estimates computed by APLS algorithms are good approximations to the solutions of AQP algorithms. In addition, we show that the performances of APLS algorithms scale to the dimensions of modern data sets. We discuss the application of our algorithms to data from European ecotypes of *Arabidopsis thaliana*, for which individual genomic a geographic data are available (Horton et al., 2012).

## 2. New methods

In this section we present two new algorithms for estimating individual admixture coefficients and ancestral genotype frequencies assuming *K* ancestral populations. In addition to genotypes, the new algorithms require individual geographic coordinates of sampled individuals.

### Q and G-matrices

Consider a genotypic matrix, **Y**, recording data for *n* individuals at *L* polymorphic loci for a *p*-ploid species (common values for *p* are *p* = 1, 2). For autosomal SNPs in a diploid organism, the genotype at locus ℓ is an integer number, 0, 1 or 2, corresponding to the number of reference alleles at this locus. In our algorithms, disjunctive forms are used to encode each genotypic value as the indicator of a heterozygote or a homozygote locus /citepFrichot2014. For a diploid organism each genotypic value 0, 1, 2 is encoded as 100, 010 and 001. For *p*-ploid organisms, there are (*p* + 1) possible genotypic values at each locus, and each value corresponds to a unique disjunctive form. While our focus is on SNPs, the algorithms presented in this section extend to multi-allelic loci without loss of generality. Moreover, the method can be easily extended to genotype likelihoods by using the likelihood to encode each genotypic value (Korneliussen et al., 2014)

Our algorithms provide statistical estimates for the matrix **Q** *∈*ℝ*K*×*n* which contains the admixture coefficients, **Q**_*i,k*_, for each sampled individual, *i*, and each ancestral population, *k*. The algorithms also provide estimates for the matrix **G** *∈* ℝ ^(*p*+1)*L*×*K*^, for which the entries, **G**_(*p*+1)*.ℓ*+*j,k*_, correspond to the frequency of genotype *j* at locus ℓ in population *k*. Obviously, the *Q* and *G*-matrices must satisfy the following set of probabilistic constraints

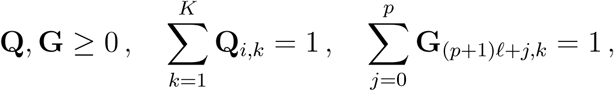

for all *i, k* and ℓ. Using disjunctive forms and the law of total probability, estimates of **Q** and **G** can be obtained by factorizing the genotypic matrix as follows **Y**=**Q G**^*T*^ (Frichot et al., 2014). Thus the inference problem can be solved by using constrained nonnegative matrix factorization methods (Lee et al., 1999; Cichocki et al., 2009). In the sequel, we shall use the notations Δ_*Q*_ and Δ_*G*_ to represent the sets of probabilistic constraints put on the **Q** and **G** matrices respectively.

### Geographic weighting

Geography is introduced in the matrix factorization problem by using weights for each pair of sampled individuals. The weights impose regularity constraints on ancestry estimates over geographic space. The definition of geographic weights is based on the spatial coordinates of the sampling sites, (*x*_*i*_). Samples close to each other are given more weight than samples that are far apart. The computation of the weights starts with building a complete graph from the sampling sites. Then the weight matrix is defined as follows

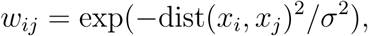

where dist(*x*_*i*_*, x*_*j*_) denotes the geodesic distance between sites *x*_*i*_ and *x*_*j*_, and *σ* is a range parameter.

Next, we introduce the *Laplacian matrix* associated with the geographic weight matrix, −**W**. The Laplacian matrix is defined as **Λ** = **D***-* **W** where **D** is a diagonal matrix with entries 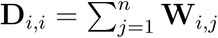, for *i* = 1*, …, n* (Belkin et al.). Elementary matrix algebra shows that (Cai et al., 2011)

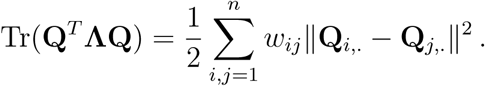

In our approach, assuming that geographically close individuals are more likely to share ancestry than individuals at distant sites is thus equivalent to minimizing the quadratic form C(**Q**) = Tr(**Q**^*T*^ **ΛQ**) while estimating the matrix **Q**.

### Least-squares optimization problems

Estimating the matrices **Q** and **G** from the observed genotypic matrix **Y** is performed through solving an optimization problem defined as follows (Caye et al., 2016)

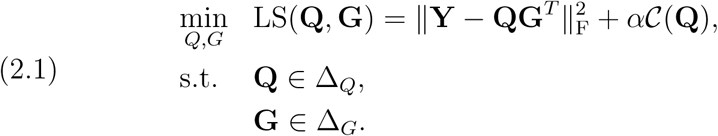

The notation ||**M**|| _F_ denotes the Frobenius norm of a matrix, **M**. The regularization parameter *α* controls the regularity of ancestry estimates over geographic space. Large values of *α* imply that ancestry coefficients have similar values for nearby individuals, whereas small values ignore spatial autocorrelation in observed allele frequencies.

### The Alternating Quadratic Programming (AQP) method

Because the polyedrons Δ_*Q*_ and Δ_*G*_ are convex sets and the LS function is convex with respect to each variable **Q** or **G** when the other one is fixed, the problem (2.1) is amenable to the application of block coordinate descent (Bertsekas, 1995). The APQ algorithm starts from initial values for the *G* and *Q*-matrices, and alternates two steps. The first step computes the matrix **G** while **Q** is kept fixed, and the second step permutates the roles of **G** and **Q**. Let us assume that **Q** is fixed and write **G** in a vectorial form, *g* = vec(**G**) ∈ R^*K*(*p*+1)*L*^. The first step of the algorithm actually solves the quadratic programming subproblem,

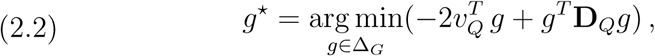

where **D**_*Q*_ = **I**_(*p*+1)*L*_ ⊗ **Q**^*T*^ **Q** and *v*_*Q*_ = vec(**Q**^*T*^ **Y**). Here, ⊗ denotes the Kronecker product and **I**_*d*_ is the identity matrix with *d* dimensions. The block structure of the matrix **D**_*Q*_ allows us to decompose the subproblem (2.2) into *L* independent quadratic programming problems with *K*(*p* + 1) variables. Now, consider that **G** is the value obtained after the first step of the algorithm, and write **Q** in a vectorial form, *q* = vec(**Q**) ∈ ℝ^*nK*^. The second step solves the following quadratic programming subproblem. Find

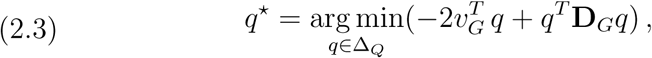

where **D**_*G*_ = **I**_*n*_⊗**G**^*T*^ **G** + *a***Λ**⊗**I**_*K*_ and *v*_*G*_ = vec(**G**^*T*^ **Y**^*T*^). Unlike sub-problem (2.2), sub-problem (2.3) can not be decomposed into smaller problems. Thus, the computation of the second step of the AQP algorithm implies to solve a quadratic programming problem with *nK* variables which can be problematic for large samples (*n* is the sample size). The AQP algorithm is described in details in Appendix A.1. For AQP, we have the following convergence result.

#### Theorem 2.1.

*The AQP algorithm converges to a critical point of problem* (2.1).

Proof. The quadratic convex functions defined in subproblems (2.2) and (2.3) have finite lower bounds. The convex sets Δ_*Q*_ and Δ_*G*_ are compact non-empty sets. Thus the sequence generated by the AQP algorithm is well-defined, and has limit points. According to Corollary 2 of Grippo et al. (2000), we conclude that the AQP algorithm converges to a critical point of problem (2.1).

### Alternating Projected Least-Squares (APLS)

In this paragraph, we introduce an APLS estimation algorithm which approximates the solution of problem (2.1), and reduces the complexity of the AQP algorithm. The APLS algorithm starts from initial values of the *G* and *Q*-matrices, and alternates two steps. The matrix **G** is computed while **Q** is kept fixed, and *vice versa*. Assume that the matrix **Q** is known. The first step of the APLS algorithm solves the following optimization problem. Find

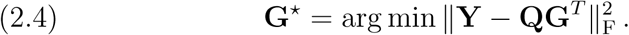

This operation can be done by considering (*p* + 1)*L* (the number of columns of **Y**) independent optimization problems running in parallel. The operation is followed by a projection of **G**^*\**^ on the polyedron of constraints, Δ_*G*_. For the second step, assume that **G** is set to the value obtained after the first step is completed. We compute the eigenvectors, **U**, of the Laplacian matrix, and we define the diagonal matrix **Δ** formed by the eigenvalues of **Λ** (The eigenvalues of **Λ** are non-negative real numbers). According to the spectral theorem, we have

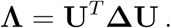

After this operation, we project the data matrix **Y** on the basis of eigenvectors as follows

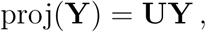

and, for each individual, we solve the following optimization problem

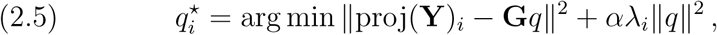

where proj(**Y**)_*i*_ is the *i*th row of the projected data matrix, proj(**Y**), and *Λ*_*i*_ is the *i*th eigenvalue of **Λ**. The solutions, 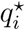, are then concatenated into a matrix, conc(*q*), and **Q** is defined as the projection of the matrix **U**^*T*^ conc(*q*) on the polyedron Δ_*Q*_. The complexity of step (2.5) grows linearly with *n*, the number of individuals. While the theoretical convergence properties of AQP algorithms are lost for APLS algorithms, the APLS algorithms are expected to be good approximations of AQP algorithms. The APLS algorithm is described in details in Appendix A.2.

### Choice of hyper-parameters

In ancestry estimation programs, a number of practices have evolved in order to set the model hyper-parameters. Those practices rely on heuristics or empirical rules for determining the prior parameters. For example, the program `structure` implements weakly informative prior distributions for ancestry proportions (Wang, 2017), the program `admixture` has a set of regularization parameters that encourages shrinkage and aggressive parsimony on ancestry estimates (Alexander et al., 2011), and so does the Bayesian version TESS 2.3 (Durand et al., 2009). Choosing the number of ancestral populations is based on cross-validation methods or information theoretic measures. Our model has three hyper-parameters: the number of factors, *K*, the penalty constant, *α*, and the range parameter, *σ*. Determining those constants is notoriously difficult and can be costly in applications. In order to reduce the computational burden, the hyper-parameters *α* and *σ* are set as user-defined options. This option allows an advanced user to explore different values with cross-validation or with her own heuristics. Less advanced users could use the default values of the hyper-parameters evaluated in our simulation study.

### Range parameter

Testing correlations between genetic and geographic data has a long tradition in population genetics. Popular approaches are based on Mantel tests (Mantel, 1967) and spatial autocorrelation measures (Hardy et al., 1999; Epperson et al., 1996). Prior to the application of our spatial ancestry estimation program, we investigated biologically relevant values for the range parameter by using spatial variograms (Cressie, 1993). The variogram was extended to genotypic data as follows

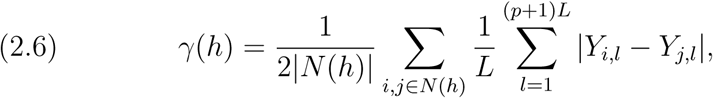

where *N* (*h*) is defined as the set of individuals separated by geographic distance *h*. Visualizing the variogram provides useful information on the level of spatial autocorrelation in the data, and yields empirical estimates of the range parameter. More naive estimates such as an average geodesic distance computed over a fraction of neighboring sites in the sample also performed well in simulations, and they are also proposed to the program users.

### Regularization parameter

A default value for the regularization parameter *α* was set so that the weights for the loss function and for the penalty term 𝒞 (**Q**) are of similar order. We proposed to divide each term by its maximum value. This amounts to consider *α* equal to *L/Λ*_max_, where *Λ*_max_ is the largest eigenvalue of the Laplacian matrix (The Laplacian matrix has nonnegative eigenvalues).

### Number of factors

The number of ancestral populations, *K*, can be evaluated by using a cross-validation technique based on imputation of masked genotypes (Wold, 1978; Eastment et al., 1982; Alexander et al., 2011; Frichot et al., 2014). The cross-validation procedure partitions the genotypic matrix entries into a learning set and a test set in which 5% of all genotypes are tagged as masked entries. The genotype probabilities for the masked entries are predicted from the factor estimates obtained from unmasked entries. Then, the error between the predicted and truly observed genotype frequencies is computed, and smaller values of that criterion indicate better choices.

### Comparison with tess3

The algorithm implemented in a previous version of `tess3` also provides another approximation of the solution of problem (2.1). The `tess3` algorithm first computes a Cholesky decomposition of the Laplacian matrix. Then, by a change of variables, the least-squares problem is transformed into a sparse nonnegative matrix factorization problem (Caye et al., 2016). Solving the sparse nonnegative matrix factorization problem relies on the application of existing methods (Kim et al., 2011; Frichot et al., 2014). The methods implemented in `tess3` have an algorithmic complexity that increases linearly with the number of loci and the number of clusters. They lead to estimates that accurately reproduce those of the Monte Carlo algorithms implemented in the Bayesian method `tess` 2.3 (Caye et al., 2016). Like for the AQP method, the `tess3` algorithms have an algorithmic complexity that increases quadratically with the sample size.

### Ancestral population differentiation statistics and local adaptation scans

Assuming *K* ancestral populations, the *Q* and *G*-matrices obtained from the AQP and from the APLS algorithms were used to compute single-locus estimates of a population differentiation statistic similar to *F*_ST_, as follows

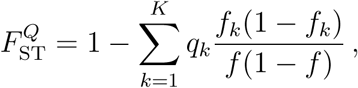

where *q*_*k*_ is the average of ancestry coefficients over sampled individuals,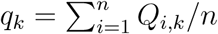, for the cluster *k*, *f*_*k*_ is the ancestral allele frequency in population *k* at the locus of interest,

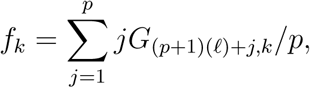

and 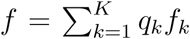 (Martins et al., 2016). For a particular locus, the formula for 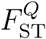 corresponds to the proportion of the genetic variation (or variance) in ancestral allele frequency that can be explained by latent population structure

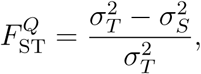

Where 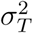 is the total variance and 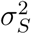 is the error variance (Weir, 1996). Following ANOVA theory, the 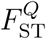 statistics were used to perform statistical tests of neutrality at each locus, by comparing the observed values to their expectations from the genome-wide background. The test was based on the squared *z*-score statistic, 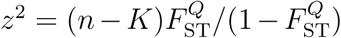 for which a chi-squared distribution with *K* − 1 degrees of freedom was assumed under the null-hypothesis. To avoid an increased number of false positive tests, we adopted an empirical null-hypothesis testing approach that recalibrates the null-hypothesis for the background levels of population differentiation expected at selectively neutral SNPs (Efron, 2004). The calibration of the null-hypothesis was achieved by using genomic control to adjust the test statistics (Devlin et al., 1999; François et al., 2016). After recalibration of the null-hypothesis, the control of the false discovery rate was achieved by using the Benjamini-Hochberg algorithm (Benjamini et al.).

`R` *package.* We implemented the AQP and APLS algorithms and improved graphical tools in the `R` package `tess3r`, available from Github and submitted to the Comprehensive R Archive Network (R Core Team, 2016).

## 3. Simulated and real data sets

### Coalescent simulations

We used the computer program `ms` to per-form coalescent simulations of neutral and outlier SNPs under spatial models of admixture (Hudson, 2002). Two ancestral populations were created from the simulation of Wright’s two-island models. The simulated data sets contained admixed genotypes for *n* individuals for which the admixture proportions varied continuously along a longitudinal gradient (Durand et al., 2009; François et al., 2010). In those scenarios, individuals at each extreme of the geographic range were representative of their population of origin, while individuals at the center of the range shared intermediate levels of ancestry in the two ancestral populations (Caye et al., 2016). For those simulations, the *Q* matrix, **Q**_0_, was entirely described by the location of the sampled individuals.

Neutrally evolving ancestral chromosomal segments were generated by simulating DNA sequences with an effective population size *N*_0_ = 10^6^ for each ancestral population. The mutation rate per bp and generation was set to *μ* = 0.25 × 10^*-*7^ the recombination rate per generation was set to *r* = 0.25 × 10^*-*8^, and the parameter *m* was set to obtain neutral levels of *F*_ST_ ranging between values of 0.005 and 0.10. The number of base pairs for each DNA sequence was varied between 10k to 300k to obtain numbers of polymorphic loci ranging between 1k and 200k after filtering out SNPs with minor allele frequency lower than 5%. To create SNPs with values in the tail of the empirical distribution of *F*_ST_, additional ancestral chromosomal segments were generated by simulating DNA sequences with a migration rate *m*_*s*_ lower than *m*. The simulations reproduced the reduced levels of diversity and the increased levels of differentiation expected under hard selective sweeps occurring at one particular chromosomal segment in ancestral populations (Martins et al., 2016). For each simulation, the sample size was varied in the range *n* = 50-700.

We compared the AQP and APLS algorithm estimates with those obtained with the `tess3` algorithm. Each program was run 5 times on the same simulated data. Using *K* = 2 ancestral populations, we computed the root mean squared error (RMSE) between the estimated and known values of the *Q*-matrix, and between the estimated and known values of the *G*-matrix. To evaluate the benefit of spatial algorithms, we compared the statistical errors of APLS algorithms to the errors obtained with the `snmf` method that reproduces the outputs of the `structure` program accurately (Frichot et al., 2014) (Frichot et al., 2015). To quantify the performances of neutrality tests as a function of ancestral and observed levels of *F*_ST_, we used the area under the precision-recall curve (AUC) for several values of the selection rate. Subsamples from a real data set were used to perform a runtime analysis of the AQP and APLS algorithms (*A. thaliana* data, see below). Runtimes were evaluated by using a single computer processor unit Intel Xeon 2.0 GHz.

### Application to human data

To evaluate the robustness of our approach to a situation where admixture was the consequence of large displacement rather than contact between proximal populations, we studied the case of African-American populations. This is an interesting case for which the incorporation of geographic data could potentially bias estimation of ancestry coefficients. Genotypes with minor allele frequency greater than 5% were obtained from a public release of the 1000 Genomes project phase 3 for African Americans (ASW, 61 individuals), Africans (YRI from Nigeria and LWK from Kenya, 207 individuals), and Europeans (GBR from the United Kingdom and TSI from Italy, 198 individuals) (1000 Genomes Project Consortium, 2015). A total of 6,994,677 SNPs were analyzed with geographic data corresponding to the country of origin of individual samples. We compared the estimates from the APLS algorithm applied with its default parameter settings to the results of the `snmf` program that do not make use of geographic information.

### Application to European ecotypes of Arabidopsis thaliana

We used the APLS algorithm to survey spatial population genetic structure and to investigate the molecular basis of adaptation by considering 214k SNPs from 1,095 European ecotypes of the plant species *A. thaliana* (Horton et al., 2012). The cross-validation criterion was used to evaluate the number of clusters in the sample, and a statistical analysis was performed to evaluate the range of the variogram from the data. We used `R` functions of the `tess3r` package to display interpolated admixture coefficients on a geographic map of Europe (R Core team 2016). A gene ontology enrichment analysis using the software AMIGO (Carbon et al., 2009) was performed in order to evaluate which molecular functions and biological processes might be involved in local adaptation of *A. thaliana* in Europe.

## 4. Results

### Statistical errors

We used coalescent simulations of neutral polymorphisms under spatial models of admixture to compare the statistical errors of the AQP and APLS estimates with those of the `tess3` algorithm (Caye et al., 2016). The ground truth for the *Q*-matrix (**Q**_0_) was computed from the mathematical model for admixture proportions used to generate the data. For the *G*-matrix, the ground truth matrix (**G**_0_) was computed from the empirical genotype frequencies in the two population samples before an admixture event. The root mean squared errors (RMSE) for the **Q** and **G** estimates decreased as the sample size and the number of loci increased (Figure 1). For all algorithms, the statistical errors were generally small when the number of loci was greater than 10k SNPs. Those results provided evidence that the three algorithms produced equivalent estimates of the matrices **Q**_0_ and **G**_0_. The results also provided a check that the APLS and `tess3` algorithms converged to the same estimates as those obtained after the application of the AQP algorithm, which is guaranteed to converge mathematically.

**Fig 1.**
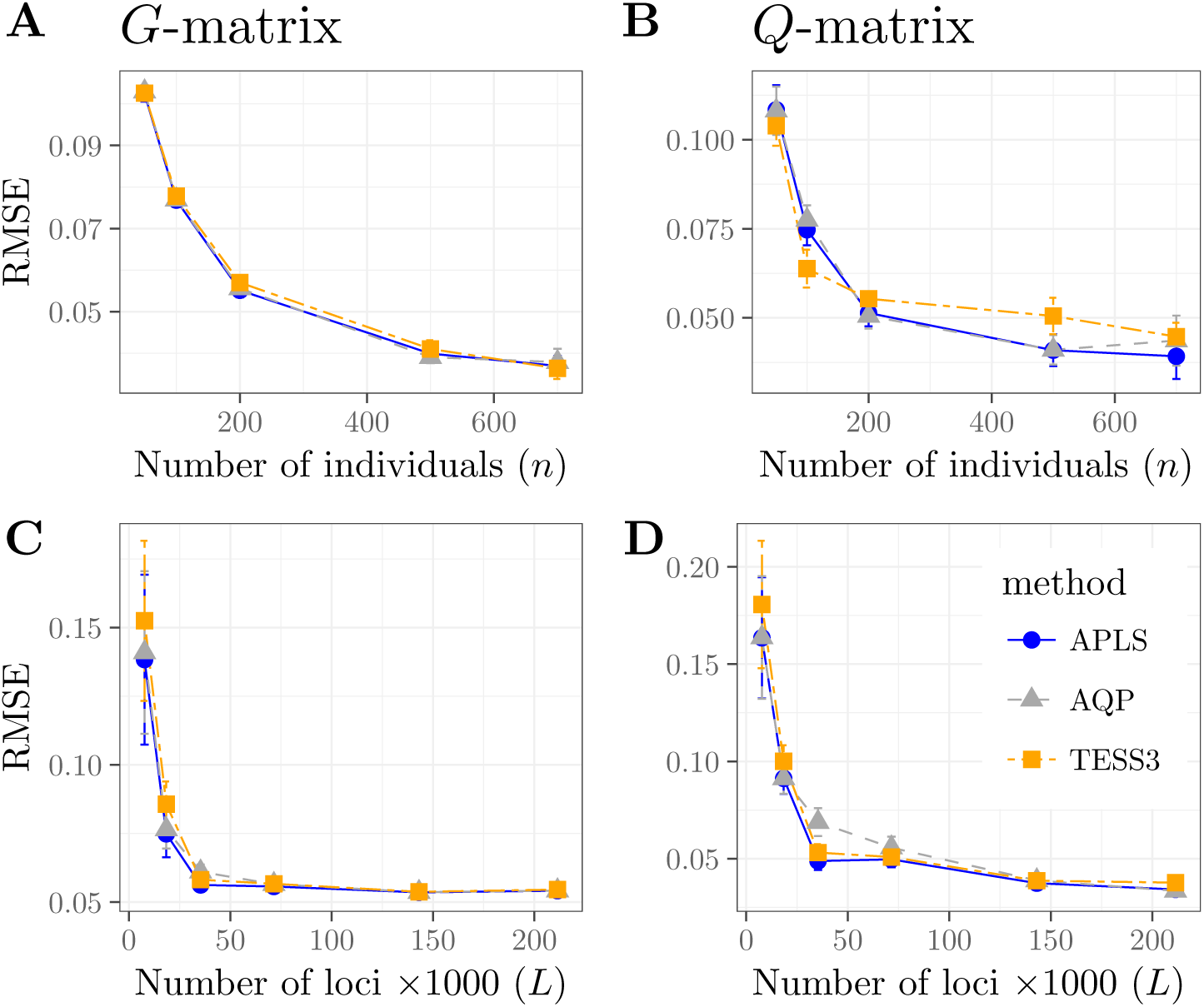
Root Mean Squared Errors (RMSEs) for the G and Q matrix estimates. Simulations of spatially admixed populations. A-B) Statistical errors for APLS, AQP and tess3 estimates as a function of the sample size, n (L 10^4^). C-D) Statistical errors for APLS, AQP and tess3 estimates as a function of the number of loci, L (n = 200).

### The benefit of including spatial information in algorithms

Using neutral coalescent simulations of spatial admixture, we compared the statistical estimates obtained from the spatial algorithm APLS and the non-spatial algorithm `snmf` (Frichot et al., 2014). For various levels of ancestral population differentiation, estimates obtained from the spatial algorithm were more accurate than for those obtained using non-spatial approaches (Figure 2). For the larger samples, much finer population structure was detected with the spatial method than with the non-spatial algorithm (Figure 2).

**Fig 2.**
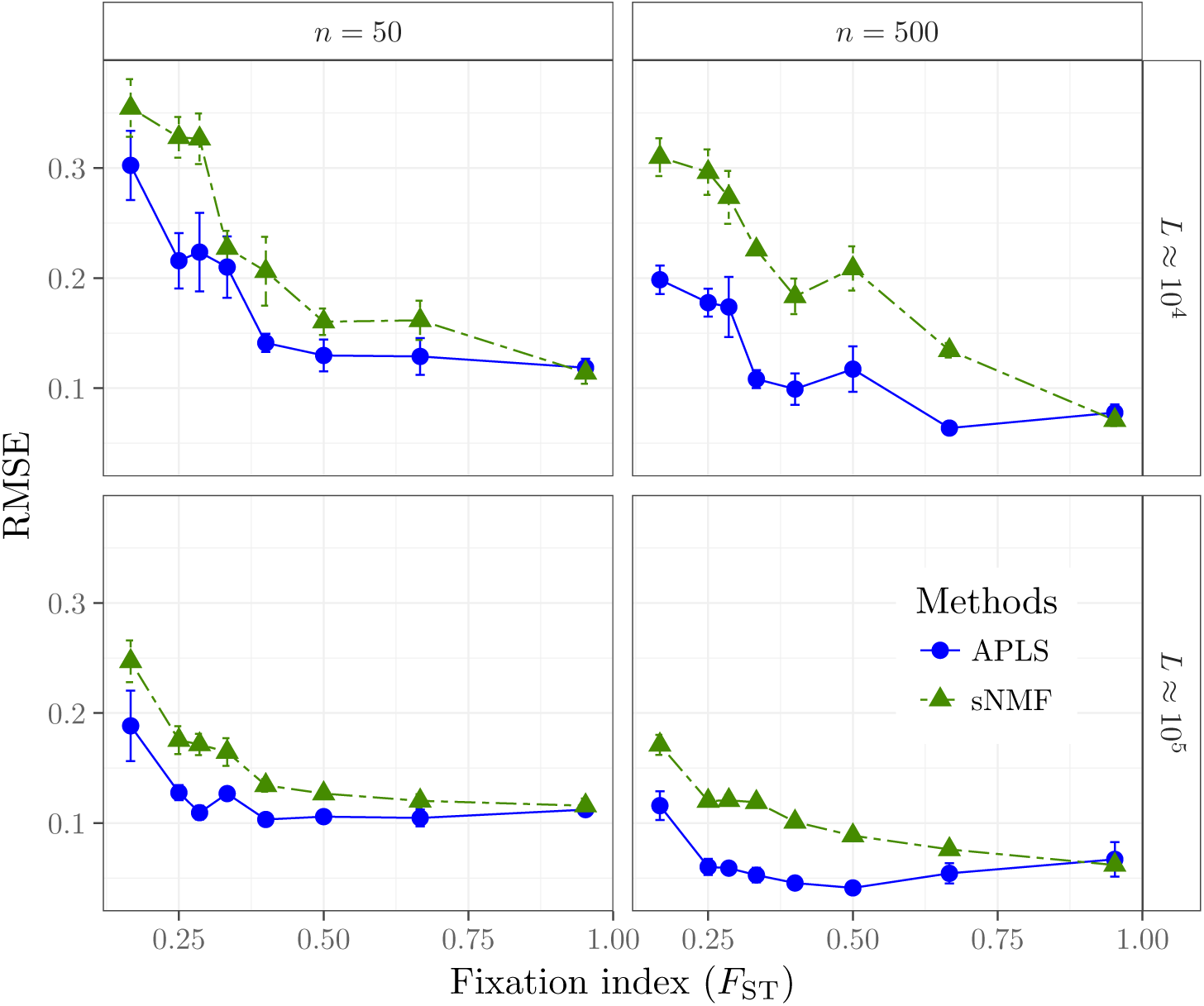
Root Mean Squared Errors (RMSEs) for the Q estimates. Simulations of spatially admixed populations for several values of fixation index (F_ST_) between ancestral populations. Ancestral populations are simulated with Wright’s two-island models and the fixation index is defined as 1/(1 + 4N_0_m) where m is the migration rate and N_0_ the effective population size. The statistical errors for sNMF and APLS are represented as a function of F_ST_.

In simulations of outlier loci, we used the area under the precision-recall curve (AUC) for quantifying the performances of tests based on the estimates of ancestry matrices, **Q** and **G**. In addition, we computed AUCs for *F*_ST_-based neutrality tests using truly ancestral genotypes. As they represented the maximum reachable values, AUCs based on truly ancestral genotypes were always higher than those obtained for tests based on reconstructed matrices. For all values of the relative selection intensity, AUCs were higher for spatial methods than for non-spatial methods (Figure 4, the relative selection intensity is the ratio of migration rates at neutral and adaptive loci). For high selection intensities, the performances of tests based on estimates of ancestry matrices were close to the optimal values reached by tests based on true ancestral frequencies. These results provided evidence that including spatial information in ancestry estimation algorithms improves the detection of signatures of hard selective sweeps having occurred in unknown ancestral populations.

**Fig 4.**
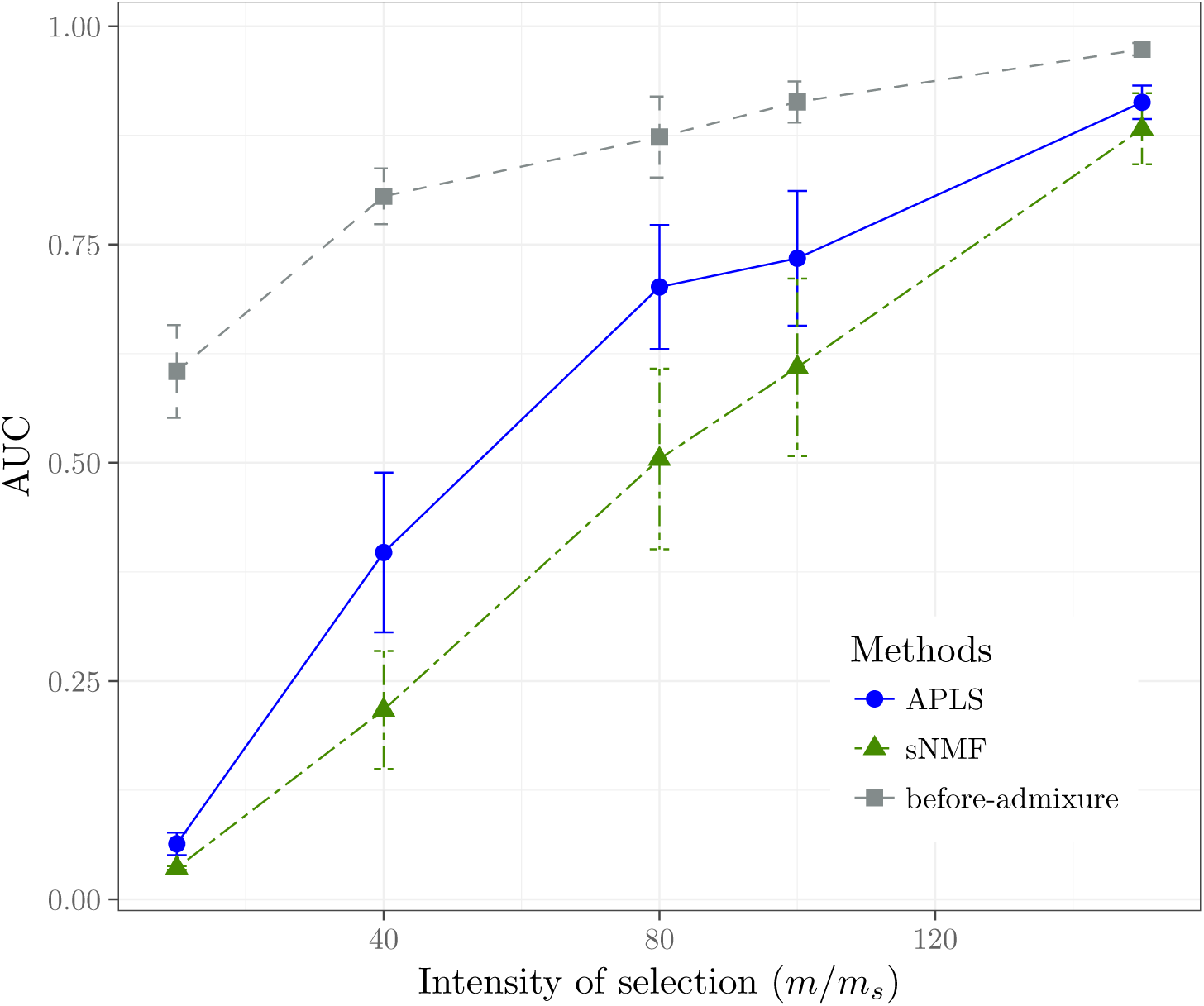
Area under the precision-recall curve (AUC). Neutrality tests applied to simulations of spatially admixed populations. AUCs for tests based on F_ST_ with the true ancestral populations, spatial ancestry estimates computed with APLS algorithms, non-spatial (structure-like) ancestry estimates computed with the snmf algorithm. The relative intensity of selection in ancestral populations, defined as the ratio m/m_s_, was varied in the range 1 - 160.

### Sensitivity of estimates to spatial measurements

Next, we used the simulated data sets to evaluate the robustness of APLS estimates to inaccurate measurements of spatial coordinates. To this aim, Gaussian noise was added to truly observed geographic coordinates by considering values of the noise-to-signal ratio ranging from 0 to 3. We computed variograms in all cases, and found that the spatial signal was removed from simulations for noise-to-signal ratios greater than two, while the signal was still observable with a noise-to-signal ratio lower than one. For all simulations, we compared the relative error of APLS *Q*-matrix estimates to those obtained from an non-spatial method (`snmf`). For small levels of uncertainty in spatial coordinates the errors of APLS estimates were lower than those of `snmf` (Figure 3). For simulations with *n* = 500 individuals and *L* = 10^5^ loci, a larger noise-to-signal ratio increased statistical errors in the *Q*-matrix estimates from the APLS algorithm. For smaller noise-to-signal ratios, RMSEs remained generally lower for the APLS algorithm than for methods without spatial coordinates. For simulations with *n* = 50 individuals and *L* = 10^4^ loci, the APLS estimates were more accurate than the non-spatial estimates. This unexpected result could be explained by subtle algorithmic differences in tested programs. To a large extent, estimates from the APLS algorithm were robust to uncertainty in spatial measurements. Standard graphical tests such as a variogram analysis can help deciding whether our spatially explicit algorithm is useful or not.

**Fig 3.**
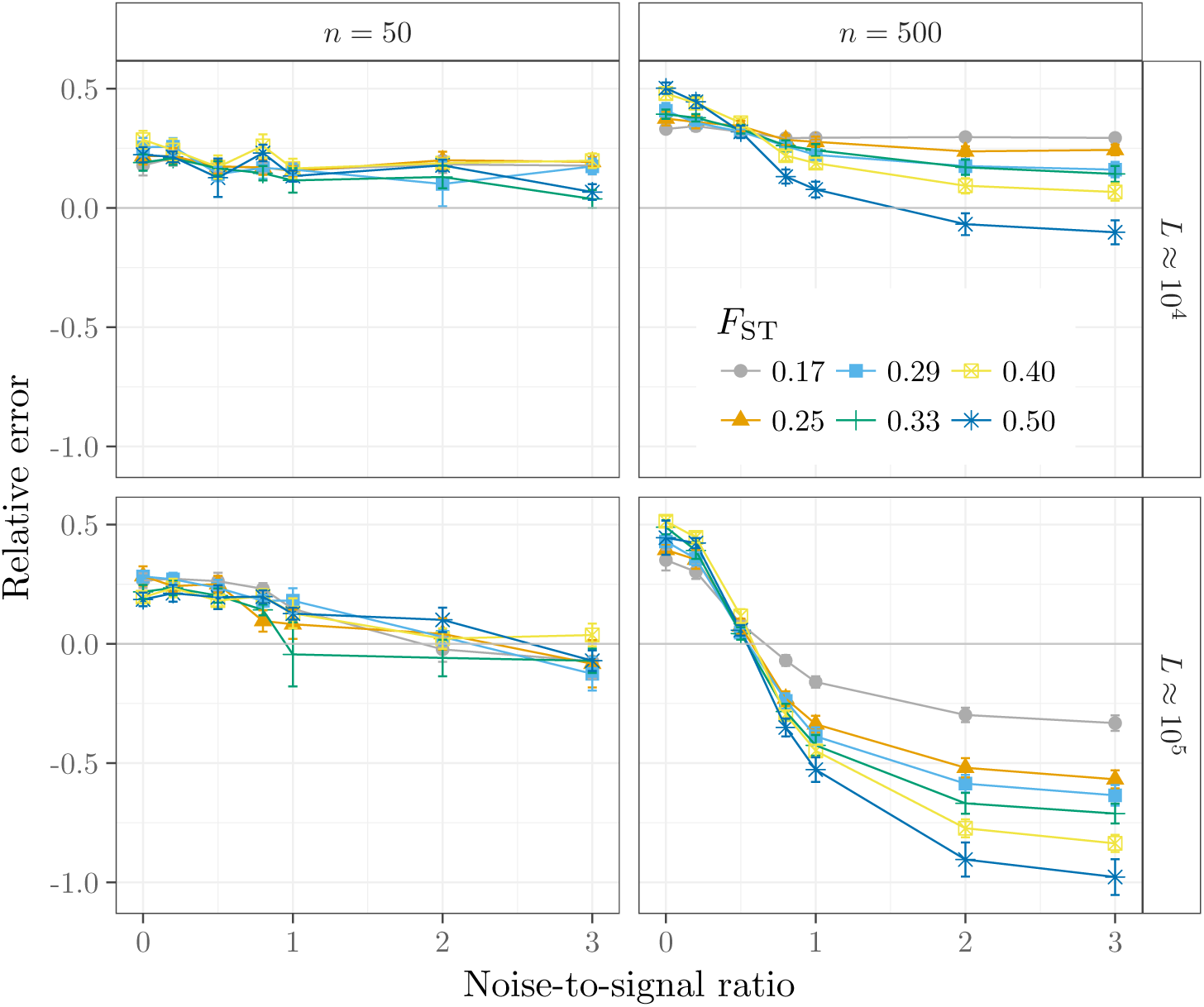
Impact of uncertainty in geographic coordinates on ancestry estimates. Relative statistical error of ancestry estimates obtained from the APLS algorithm for several levels of the noise-to-signal ratio and values of the fixation index. The snmf algorithm was considered as the reference for the non-spatial method (value 0).

### Runtime and convergence analyses

We subsampled a large SNP data set for *A. thaliana* ecotypes to compare the convergence properties and runtimes of the `tess3`, AQP, and APLS algorithms. In those experiments, we used *K* = 6 ancestral populations, and replicated 5 runs for each simulation. For *n* = 100 − 600 individuals (*L* = 50k SNPs), the APLS algorithm required more iterations (25 iterations) than the AQP algorithm (20 iterations) to converge to its solution (Figure 5). This was less than for `tess3` (30 iterations). For *L* = 10 − 200k SNPs (*n* = 150 individuals), similar results were observed. For 50k SNPs, the runtimes were significantly lower for the APLS algorithm than for the `tess3` and AQP algorithms. For *L* = 50k SNPs and *n* = 600 individuals, it took on average 1.0 min for the APLS and 100 min for the AQP algorithm to compute ancestry estimates. For `tess3`, the runtime was on average 66 min. For *L* = 100k SNPs and *n* = 150 individuals, it took on average 0.6 min (9.0 min) for the APLS (AQP) algorithm to compute ancestry estimates. For `tess3`, the runtime was on average 1.3 min. For those values of *n* and *L*, the APLS algorithm implementation ran about 2 to 100 times faster than the other algorithm implementations.

**Fig 5.**
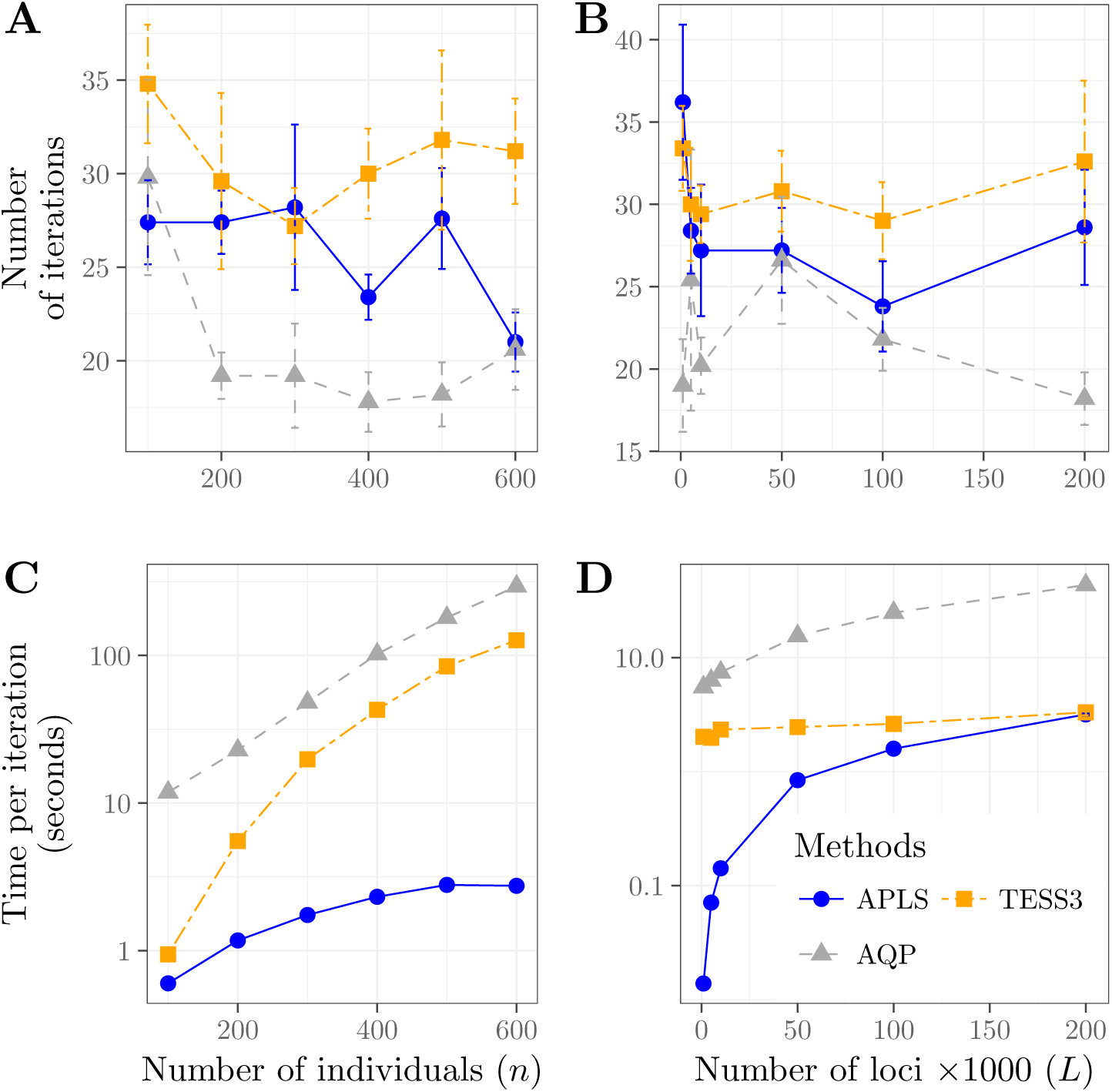
Number of iterations and runtimes for the AQP, APLS and tess3 algorithm implementations. A-B) Total number of iterations before an algorithm reached a steady solution. C-D) Runtime for a single iteration (seconds). The number of SNPs was kept fixed to L = 50k in A and C. The number of individuals was kept fixed to n = 150 in B and D.

### Human data analysis

To evaluate a case of model misspecification, we analyzed data from the 1000 Genomes project for African Americans, Africans from Nigeria and from Kenya, and Europeans from the United Kingdom and from Italy. Using the default values for the hyper-parameters, the Laplacian matrix was a block diagonal matrix where each block corresponded to one of the five populations. The spatial variogram exhibited a flat shape. For *K* = 2, the APLS estimates for the African American population were equal to 24.2% for European ancestors and 75.8% for African ancestors. The corresponding `snmf` estimates were equal to 22.4% for European ancestors and 77.6% for African ancestors. For *K* = 3, the APLS estimates for the African American population were equal to 21.4% for European ancestors, 51.8% for West African ancestors and 26.8% for East African ancestors. The corresponding `snmf` estimates were equal to 22.2% for European ancestors, 68.4% for West African ancestors and 9.4% for East African ancestors. Overall, the results obtained with our spatial method for African Americans were similar to those obtained with `snmf`. The main difference between APLS and `snmf` estimates were for African populations. For Africans, `snmf` detected two distinct genetic clusters whereas APLS detected a larger proportion of shared ancestry between Eastern and Western populations.

### Application to European ecotypes of Arabidopsis thaliana

We used the APLS algorithm to survey spatial population genetic structure and perform a genome scan for adaptive alleles in European ecotypes of the plant species *A. thaliana*. The cross validation criterion decreased rapidly from *K* = 1 to *K* = 3 clusters, indicating that there were three main ancestral groups in Europe, corresponding to geographic regions in Western Europe, Eastern and Central Europe and Northern Scandinavia. For *K* greater than four, the values of the cross validation criterion decreased in a slower way, indicating that subtle substructure resulting from complex historical isolation-by-distance processes could also be detected (Figure 6). The spatial analysis provided an approximate range of *σ* = 150km for the spatial variogram (Figure 6). Figure 7 displays the *Q*-matrix estimate interpolated on a geographic map of Europe for *K* = 6 ancestral groups. The estimated admixture coefficients provided clear evidence for the clustering of the ecotypes in spatially homogeneous genetic groups.

**Fig 6.**
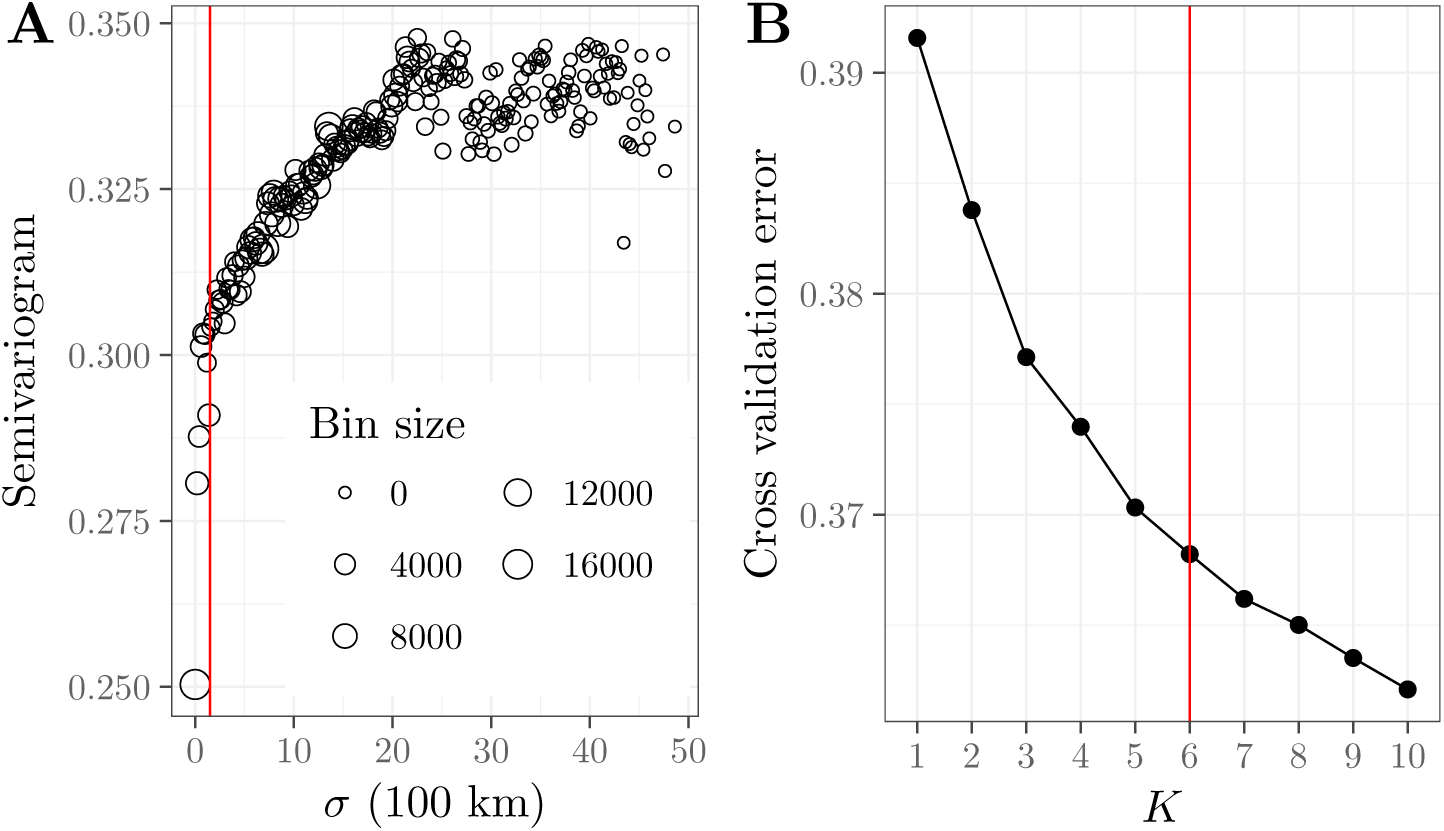
Choice of σ and K for the APLS algorithm. A) Empirical variogram for the A. thaliana data. The red vertical line shows the range value σ = 1.5. B) Cross validation error as function of the number of ancestral populations, K. The red vertical line shows the number of ancestral populations K = 6.

**Fig 7.**
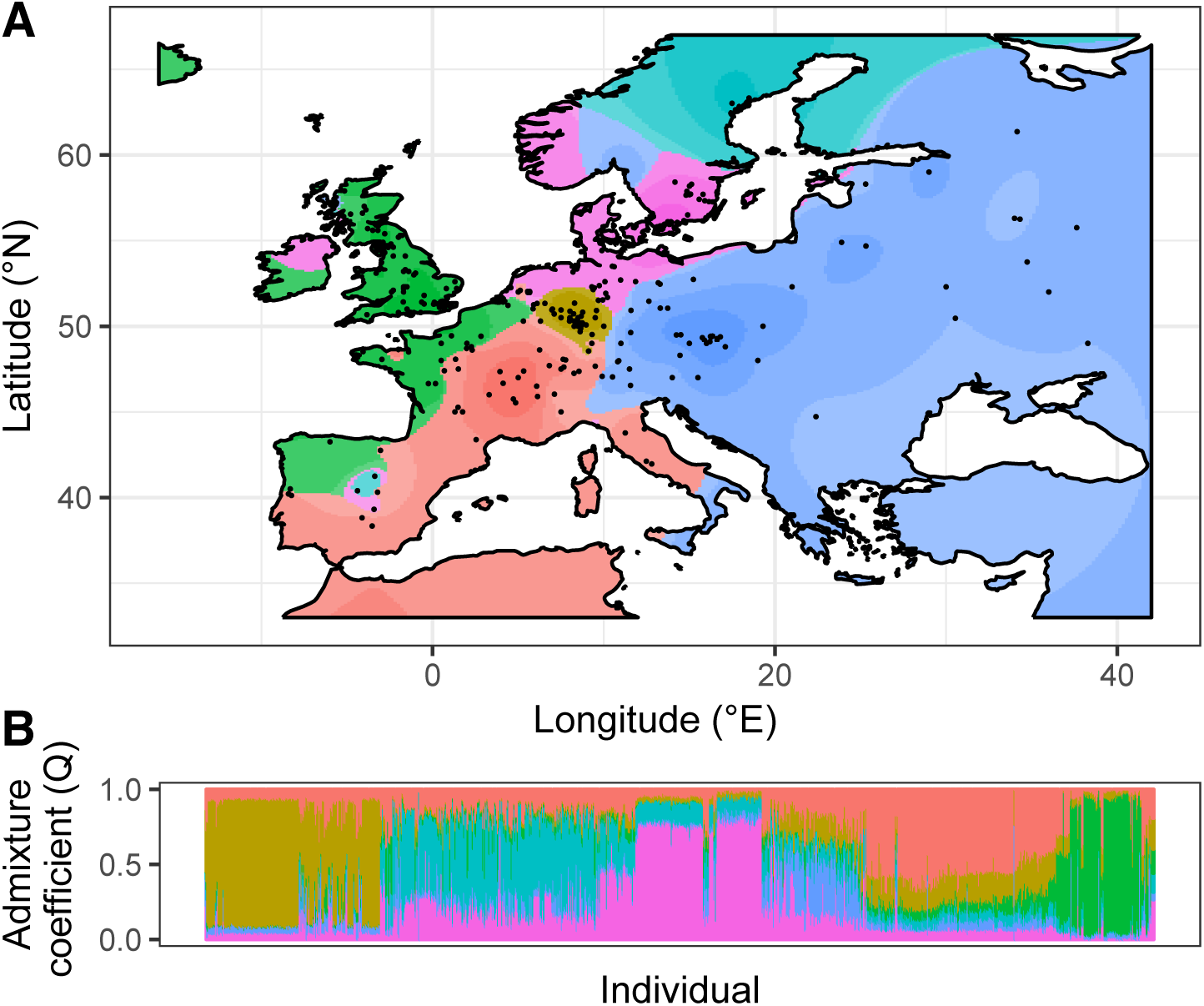
A. thaliana ancestry coeficients. Ancestry coefficient estimates computed by the APLS algorithm with K = 6 ancestral populations and σ = 1.5 for the range parameter. A) Geographic map of ancestry coefficients. B) Barplot of ancestry coefficients.

**Fig 8.**
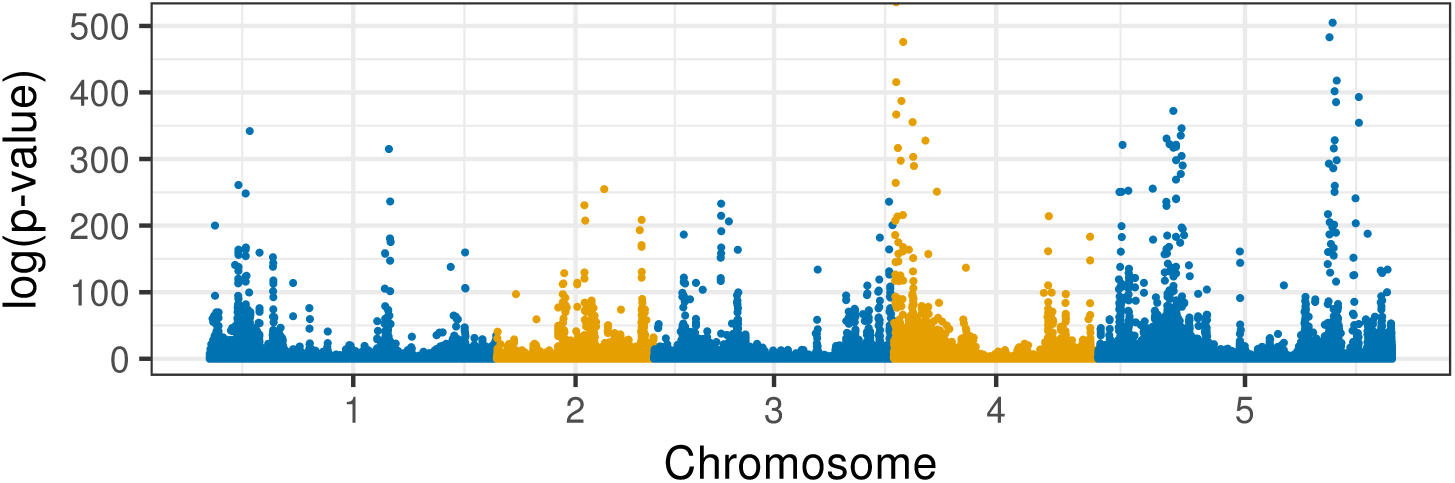
Local adaptation in European lines of. ***A***. *thaliana. Manhattan plot of* − log(*p*-value). *p*-value were computed from population structure estimated by the APLS algorithm with *K* = 6 ancestral populations and *σ* = 1.5 for the range parameter.

### Targets of selection in A. thaliana genomes

Tests based on the 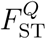 statistic were applied to the 241k SNP data set to reveal new targets of natural selection in the *A. thaliana* genome. *A. thaliana* occurs in a broad variety of habitats, and local adaptation to the environment is acknowledged to be important in shaping its genetic diversity through space (Hancock et al., 2011; Fournier-Level et al., 2011). The APLS algorithm was run on the 1,095 European lines of *A. thaliana* with *K* = 6 ancestral populations and *σ* = 1.5 for the range parameter. Using the Benjamini-Hochberg algorithm to control the FDR at the level 1%, the program produced a list of 12,701 candidate SNPs, including linked loci and representing 3% of the total number of loci. The top 100 candidates included SNPs in the flowering-related genes SHORT VEGETATIVE PHASE (SVP), COP1-interacting protein 4.1 (CIP4.1) and FRIGIDA (FRI) (*p*-values < 10^*-*300^). These genes were detected by previous scans for selection on this dataset (Horton et al., 2012). We performed a gene ontology enrichment analysis using AmiGO in order to evaluate which biological functions might be involved in local adaptation in Europe. We found a significant over-representation of genes involved in cellular processes (fold enrichment of 1.06, *p*-value equal to 0.0215 after Bonferonni correction).

## 5. Discussion

Including geographic information on sample locations in the inference of ancestral relationships among organisms is a major objective of population genetic studies (Malécot, 1948; Cavalli et al., 1994; Epperson, 2003). Assuming that geographically close individuals are more likely to share ancestry than individuals at distant sites, we introduced two new algorithms for estimating ancestry proportions using geographic information. Based on least-squares problems, the new algorithms combine matrix factorization approaches and spatial statistics to provide accurate estimates of individual ancestry coefficients and ancestral genotype frequencies. The two methods share many similarities, but they differ in the approximations they make in order to decrease algorithmic complexity. More specifically, the AQP algorithm was based on quadratic programming, whereas the APLS algorithm was based on the spectral decomposition of the Laplacian matrix. The algorithmic complexity of APLS algorithm grows linearly with the number of individuals in the sample while the method has the same statistical accuracy as more complex algorithms.

To measure the benefit of using spatial algorithms, we compared the statistical errors observed for spatial algorithms with those observed for non-spatial algorithms. The errors of spatial methods were lower than those observed with non-spatial methods, and spatial algorithms allowed the detection of more subtle population structure. In addition, we implemented neutrality tests based on the spatial estimates of the *Q* and *G*-matrices (Martins et al., 2016), and we observed that those tests had higher power to reject neutrality than those based on non-spatial approaches. Thus spatial information helped improving the detection of signatures of selective sweeps having occurred in ancestral populations prior to admixture events. We applied the neutrality tests to perform a genome scan for selection in European ecotypes of the plant species *A. thaliana*. The genome scan confirmed the evidence for selection at flowering-related genes *CIP4.1*, *FRI* and *DOG1* differentiating Fennoscandia from North-West Europe (Horton et al., 2012).

Estimation of ancestry coefficients using fast algorithms that extend non-spatial approaches – such as `structure` – has been intensively discussed during the last years (Wollstein et al., 2015). In these improvements, spatial approaches have received less attention than non-spatial approaches. In this study, we have proposed a conceptual framework for developing fast spatial ancestry estimation methods, and a suite of computer programs implements this framework in the `R` program `tess3r`. Our package provides an integrated pipeline for estimating and visualizing population genetic structure, and for scanning genomes for signature of local adaptation. The algorithmic complexity of our algorithms allows their users to analyze samples including hundreds to thousands of individuals. For example, analyzing more than one thousand *A. thaliana* genotypes, each including more than 210k SNPs, took only a few minutes using a single CPU. In addition, the algorithms have multithreaded versions that run on parallel computers by using multiple CPUs. The multithreaded algorithm, which is available from the `R` program, allows using our programs in large-scale genomic sequencing projects.

## APPENDIX A: ALGORITHMS

### Algorithm A.1.

AQP algorithm pseudo code. To solve optimization problem (2.1).

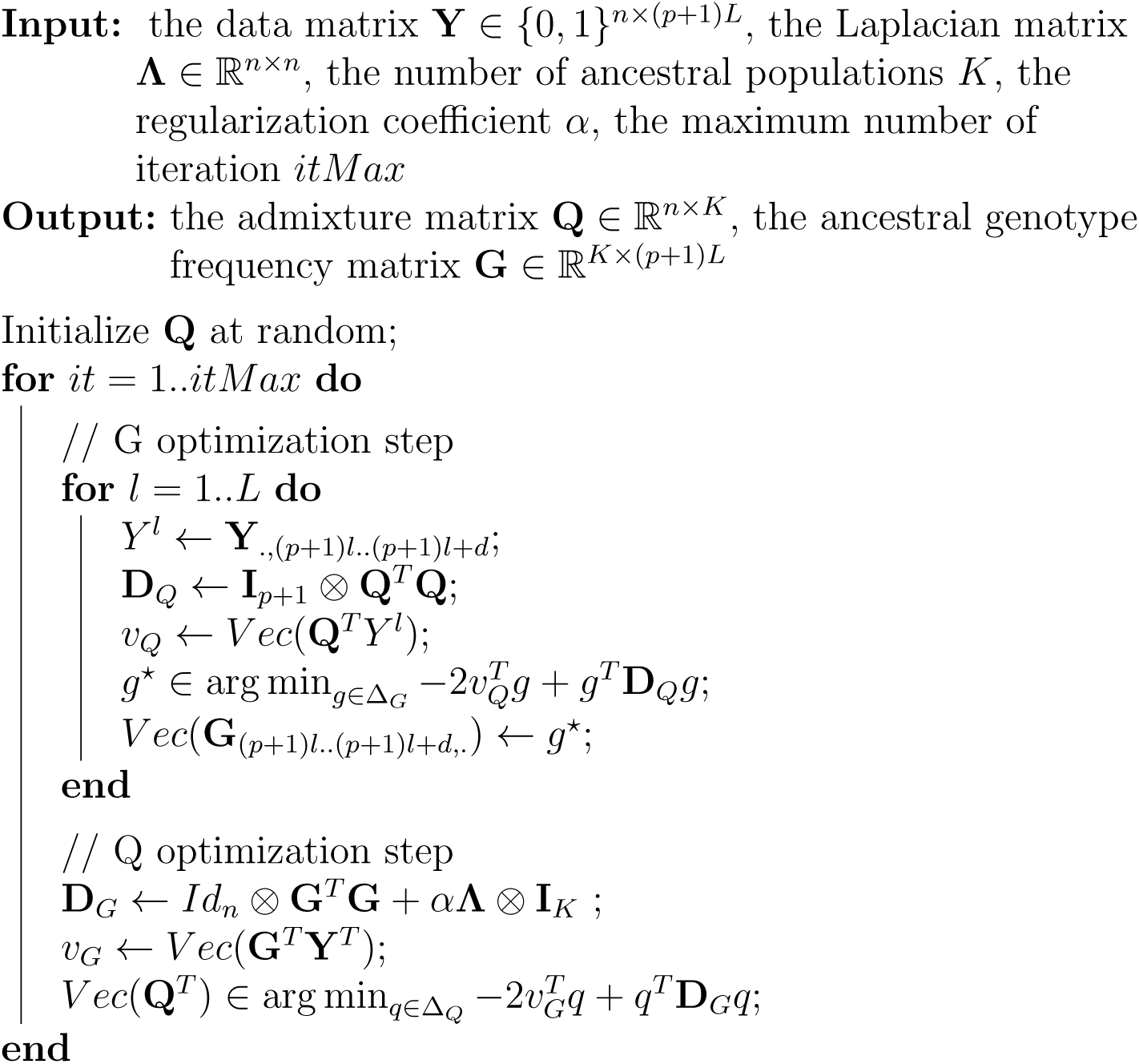

### Algorithm A.2.

APLS algorithm pseudo code. To solve the optimization problem (2.1).

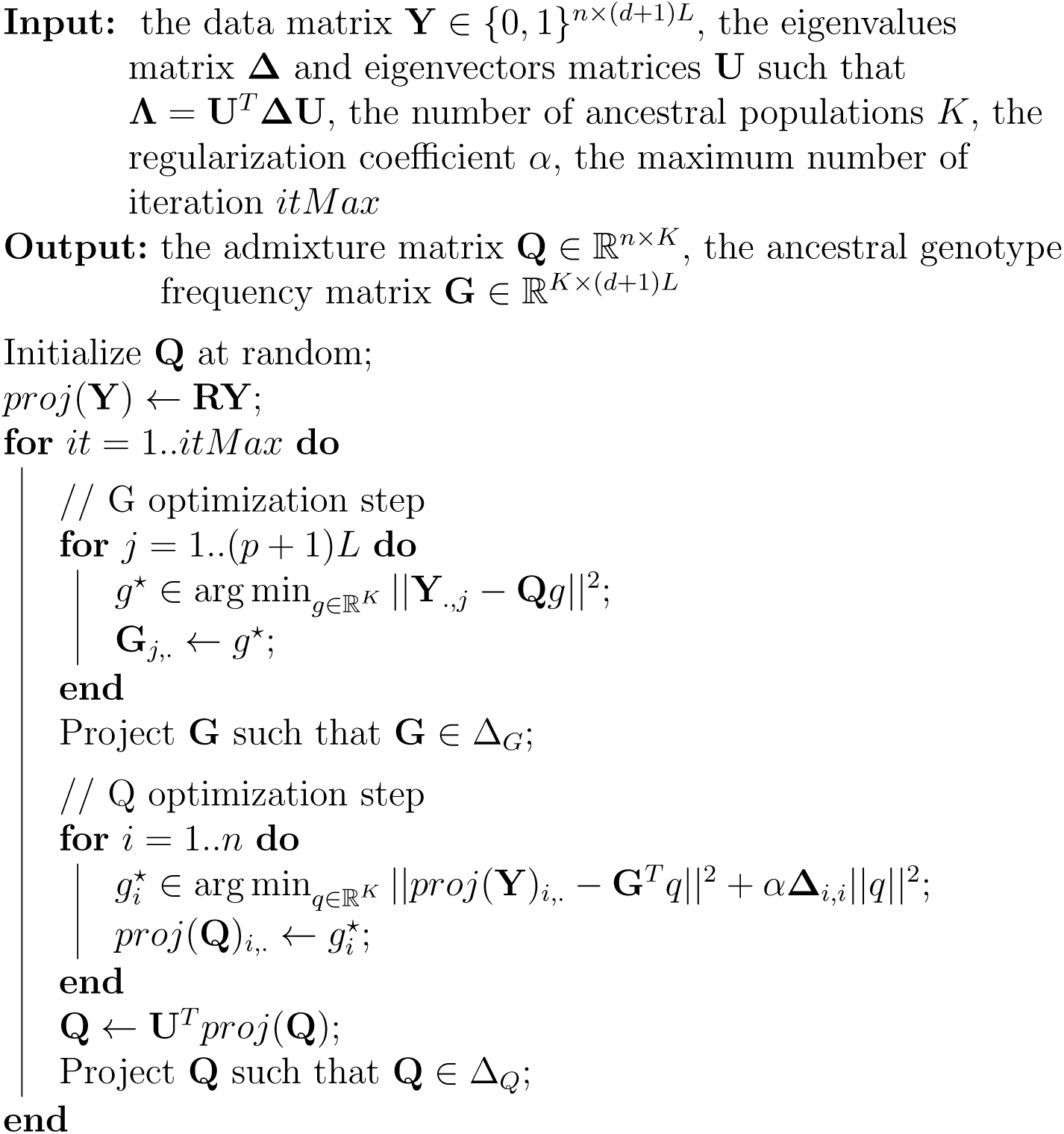

## ACKNOWLEDGEMENTS

The authors are grateful to Edoardo M. Airoldi and three anonymous reviewers for their constructive comments. This work has been partially supported by the LabEx PERSYVAL-Lab (ANR-11-LABX-0025-01) funded by the French program Investissement d’Avenir. Olivier François acknowledges support from Grenoble INP and from the Agence Nationale de la Recherche, project AFRICROP ANR-13-BSV7-0017.

